# Physiological and genetic analysis of tomato from two cultivars differing in potassium deficiency resistance

**DOI:** 10.1101/2021.05.06.442937

**Authors:** Xi Wang, Honghui Zhang, Tianlai Li, Xin Liu, Jing Jiang

## Abstract

Potassium (K) is one of the essential nutrients for tomato. Potassium deficiency will limit tomato growth and yield. So improving the low-K^+^ (LK) resistance of tomato seems important. Two tomato cultivars (JZ18 and JZ34) differing in LK resistance were obtained to analyze the plant demonstration difference under LK treatment. According to the results, JZ34 showed lower accumulation of ROS, less membrane damage and higher antioxidant enzyme activity after LK treatment. Besides, JZ34 also keeps higher K^+^/Na^+^ content, higher Ca^2+^ and Mg^2+^ content than JZ18 in both shoots and roots. Our genetic analysis revealed that the two additive-dominance-epistasis major genes plus additive-dominance polygene genetic model (E-1) was the optimum model associated with LK resistance based on root trait. The major QTL intervals were finally obtained by the bulked segregant sequencing (BSA-seq) analysis, which were 2.38 Mb at the end of chromosome 4 and 1.38 Mb at the chromosome 6. This is consistent with the analysis of the genetic model. A total of 8 genes were selected in the two candidate regions, which exhibited close related to ion and antioxidant signaling. These findings provided the inheritance pattern and foundation for further molecular mechanisms study of tomato LK resistance.

## 1. Introduction

Potassium is the most abundant cation in plant cells, and plays crucial roles in diverse physiological processes during plant growth and development, such as enzyme activation, electrical neutralization, and membrane potential maintenance (Wang and Wu 2017). While the K^+^ concentration in the soil solution may vary widely from 0.01 to 20 mM, plant cells maintain a relatively constant concentration of 80-100 mM in the cytoplasm (Rodríguez-Navarro 2000). Compared with the higher concentration of K^+^ in cells, the concentration of K^+^ in soil was lower. The roots of plants are in direct contact with the soil, so the LK signal is initially sensed by root cells, especially root epidermal cells and root hair cells. K^+^ deficiency signal is first sensed by the plasma membrane of root epidermal cells, and then transmitted to the cytoplasm, causing a series of physiological and biochemical processes in response to LK stress (Song *et al.* 2018).

In response to LK, plants affect root growth and root architecture, such as inhibiting the growth of taproot and stimulating root hair elongation (Cao *et al.* 1993; Tsay *et al.* 2011). Using some root traits as an evaluation standard, some genes or quantitative trait loci (QTLs) related to LK resistance were discovered in rice (Koyama *et al.* 2001; Lin *et al.* 2004; Pandit *et al.* 2010), wheat (Kong *et al.* 2013) and *A. thaliana* (Xu *et al.* 2006; Wang *et al.* 2010; Li *et al.* 2017; Du *et al.* 2019). Protein kinase CIPK23, interaction with the K^+^ channel AKT1, was map-based cloned by observing the different root phenotype of mutant and wild type under LK condition in *A. thaliana* (Xu *et al.* 2006). Quantitative traits loci (QTLs) for root length and root dry weight in rice were detected using a doubled haploid population under LK condition (Fang *et al.* 2015).

Tomato have a high demand for K^+^ as the major horticulture crops. LK conditions would result in the serious decline of production and quality in tomato. However, it has been rare studies in tomato to study the mapping of K^+^ deficiency resistance gene. Only few early reports showed the K^+^ utilization efficiency had low heritability, and were controlled by polygene and affected by additive effect, dominance effect and epistasis effect in tomato (Gabelman and Loughman 1987). Thus, it is important for investigations of tomato breeding of LK resistance to learn the inheritance models of tomato in response to LK stress.

In the molecular signal network response to LK stress, some signaling molecules have been proposed to be involved in, including ion, ROS signal and so on (Wang and Wu 2013). The LK stress firstly affects the plasma membrane, activates the Ca^2+^ channel, and initiates the LK signal transduction pathway (Behera *et al.* 2016). When plants suffer from LK stress, the Ca^2+^ sensor CBL can interact with the protein kinase CIPK to form a complex to activate the high-affinity K^+^ transporter or K^+^ channel, thereby responding to the LK stress (Xu *et al.* 2006; Dong *et al.* 2021). Similar to magnesium-calcium, an antagonistic relationship has also been described for magnesium-potassium (Senbayram *et al.* 2015). High levels of external K^+^ result in reduced uptake of Mg^2+^, and an effect of high Mg^2+^ on K^+^ uptake has also been reported in Arabidopsis (Fageria 2001; Ding *et al.* 2010; Mogami *et al.* 2015). Early study suggested that at least two mechanisms are involved in Mg^2+^-uptake through the plasma membrane, one of which allows for uptake of K^+^ and Ca^2+^ (Shabala and Hariadi 2005). In addition, facilitating osmotic adjustment and maintenance of high K^+^/Na^+^ ratios in the cytosol of plants is essential for salt and LK tolerance. It involves a network of transport processes that regulates uptake, extrusion through the plasma membrane, compartmentation of salts into cell vacuoles and recirculation of ions through the plant organs (Apse and Blumwald 2007; Pardo and Rubio 2011; Asins *et al.* 2013). Moreover plants will produce a large amount of reactive oxygen species (ROS) under LK stress, which are important signaling molecules in cells. Studies have shown that ROS are not only involved in LK signaling, but also are induced Ca^2+^ signaling to convey HAK5 K^+^ transporter induction (Mittler 2002; R and Schachtman 2004; Wang *et al.* 2021). How these ions and ROS signals in response to LK are transmitted in tomato is still unknown, which requires a comprehensive study to explore.

In our research, two tomato varieties with different tolerance to LK, LK-sensitive inbred line ‘JZ18’ and LK-resistant inbred line ‘JZ34’ were used to determine the difference of ion and ROS signaling in response to LK stress. Next, the length of roots values of six generations were (P_1_, P_2_, F_1_, B_1_, B_2_ and F_2_) measured under LK stress, JZ18 and JZ34 lines as parents, and major gene and polygene genetic models were acquired. The major-effect QTLs was further confirmed by BSA-seq, which led to better understand the molecular mechanism of K^+^ deficiency resistance in tomato seedlings. These results may provide a basic theory for further key signaling pathways and QTL analysis for LK resistance in tomato.

## 2. Materials and methods

### 2.1 Plant materials and growth condition

Through observation of 9 tomato materials with different stem diameter, root activity, root dry weight, root fresh weight and yield under LK condition in the 2010, two lines were selected from them, JZ18 (P_1_), with LK susceptibility, and JZ34 (P_2_), with high resistance to LK (Zhao *et al.* 2018). In autumn 2017, JZ18 (P_1_) and JZ34 (P_2_) as the parents were crossed to construct F_1_ populations in experimental field of Shenyang Agricultural University. In autumn 2018, the F_2_ generation was produced by strict self-pollination of F_1_ and BC_1_P_1_ and BC_1_P_2_ generations were obtained using F_1_ and two parents to backcross.

The seed of six generations were sterilized in culture dish to germinate and were transferred in plug plate after 3 days. All the seedlings were provided with the same growth conditions. The LK condition was generated by hydroponics method (Zhao *et al.* 2018). The K^+^ concentration of control solution was 4 mM and LK solution was 0.5 mM. After LK stress for 21 days, the samples of the tomato seedlings were taken to obtain the pictures and determined the chlorophyll, relative water content (RWC), biomass, and proline contents. Three biological replicates were conducted for each treatment.

### 2.2 Observation and determination of root morphology

The root materials were obtained after LK stress treatment for 7 days. The root traits of the tomato seedlings were scanned by the Epson Scan 2 and analyzed WinRHIZO software, including root length, root area and root fork. The phenotypic data of root length, including maxinum, mininum, mean, standard deviation and variance, were obtained by Excel 2010. The CV (%) was evaluated by formulas, (CV: coefficient of variation, σ: standard deviation, μ: average).

### 2.3 ROS and ion content measurement

The JZ18 and JZ34 plants, after LK stress for 0, 1, 3 and 7 days and these plants were used for the measurements. Three biological replicates were conducted for each treatment. The content of H_2_O_2_ and O^2 −^ was detected using the Micro Hydrogen Peroxide(H_2_O_2_) Assay Kit (Solarbio Science, China). The malondialdehyde (MDA) content, the activities of super oxide dismutase (SOD), peroxidase (POD), catalase (CAT), and ascorbate peroxidase (APX) and the K^+^, Na^+^, Ca^2+^ and Mg^2+^ content were determined (Zhao *et al.* 2017).

### 2.4 Joint segregation analysis

The mix major gene plus polygene genetic models were obtained by the joint segregation analysis method using the phenotypic values of six generations (Gai and Wang 1998). The 24 types inheritance models were classified for five groups, including one major gene model (A), two major genes model (B), polygene model (C), one major gene plus polygene model (D) and two major genes plus polygene model (E). The two models with smallest values of AIC were selected as candidate models. A series of goodness-of-fit test, including the homogeneity test (U_2_^1^, U_2_^2^ and U_2_^3^), Smirnov test (nW^2^) and Kolmogorov test (D_n_), were used to estimate the candidate models. The model with the smallest number of significance was chosen as the best-fit model. Finally, the first-and second-order parameters of the best model were acquired.

### 2.5 BSA-seq and linkage mapping analysis

For bulked segregant analysis (BSA), two extreme pools were selected from the F_2_ population (650 individuals): a LK resistance pool (R-pool, 28 F_2_ individuals showing 5000 cm root length) and a LK sensitive pool (S-pool, 28 F_2_ individuals showing 800 cm root length). Total DNA was extracted from the parental lines and extreme pools, and used for library construction for short-insert sequencing. Qualified DNA samples are broken into 400bp fragments by the fragmentation kit for library construction. The DNA fragments are subjected to end repair, polyA tailing, sequencing adapters, purification, PCR amplification and other steps to complete the entire library preparation. After the library is constructed, use Qubit2.0 for preliminary quantification, and then use the Agilent BioAnalyzer 2100 to detect the length of the insert in the library. After the length meets the expectation, use qPCR to accurately quantify the effective concentration of the library to ensure the quality of the library. After the library is qualified, it enters the superior sequencing stage. The sequencing platform is Illumina Hiseq 4000, and the sequencing mode is PE150. And then, we will perform quality control on the offline data, and get CleanData after removing low-quality sequences and sequencing adapter sequences. Next, compare these CleanData data with the reference genome (https://www.solgenomics.net/organism/Solanum_lycopersicum/genome), and perform SNP and InDel detection and annotation based on the comparison results. Next, calculate the SNP-index and the difference of the offspring mixed pools, select the regions with very significant differences in SNP-index of the two offspring mixed pools, and locate the target trait regions on the *Solanum lycopersicums*’ chromosomes.

## 3. Results

### 3.1 Phenotypic characterization of LK tolerance in JZ18 and JZ34

The two self-bred lines of tomato, JZ18 (LK susceptible lines), JZ34 (LK resistant lines), grown on normal conditions were transferred to LK conditions for another 7 days. The JZ18 displayed root growth inhibition under LK conditions for 7 days(**Fig. 1** A). And JZ18 plants had the smaller whole root area, shorter root length and lower root fork under LK stress. (**Fig. 1** B-D). However, JZ34 had a more stable root system under LK stress. These results demonstrate that LK stress restrained the root growth of the JZ18 plants, while almost no effect on JZ34 plants(**Fig. 1** E).

**Fig. 1.**
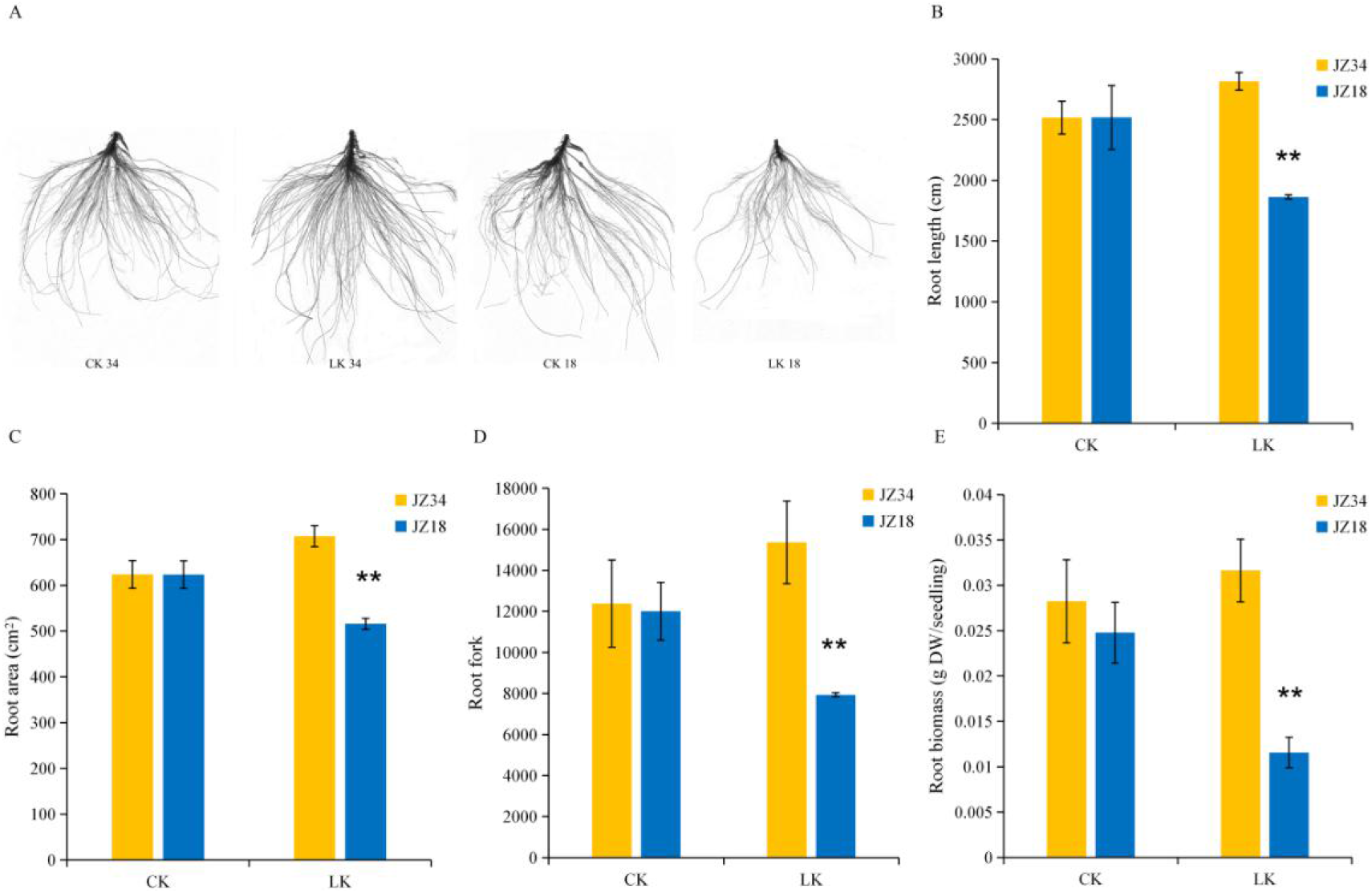
Roots phenotype of JZ18 and JZ34 under control K^+^ (4 mM) and LK (0.5 mM) stress conditions for 7 days. (A) Root phenotype, (B) Root length, (C) Root area, (D) Root fork and (E) Root biomass in JZ18 and JZ34 plants under control and LK stress conditions for 7 days. The presented data are the means ± SE of three independent experiments (n=12). Asterisks indicate significant difference between JZ18 and JZ34 plants (*P < 0.05, **P < 0.01).

In addition, when JZ18 and JZ34 plants were transferred to LK conditions for 21 days, the leaves turned yellow-green color in the JZ18, which is a typical K^+^ deficiency phenotype, while the leaves growth of JZ34 remained normal green color (**Fig. 2** A). As showed in **Fig. 2** B-C, pigment measurement results indicated that the contents of chlorophyll a, chlorophyll b and carotenoids in JZ18 and JZ34 were significantly (P value < 0.05) lower under LK stress compared with those in control (CK), while the degree of decrease in JZ34 was smaller than JZ18 (**Fig. 2** B-D). Similar results were observed in relative water content and shoot biomass (**Fig. 2** E-F). After exposure to LK stress, the proline content of JZ34 plants was increased significantly (**Fig. 2** G). These results suggesting that the LK tolerance of JZ34 plants was higher than JZ18 plants through maintaining normal root and leaf growth.

**Fig. 2.**
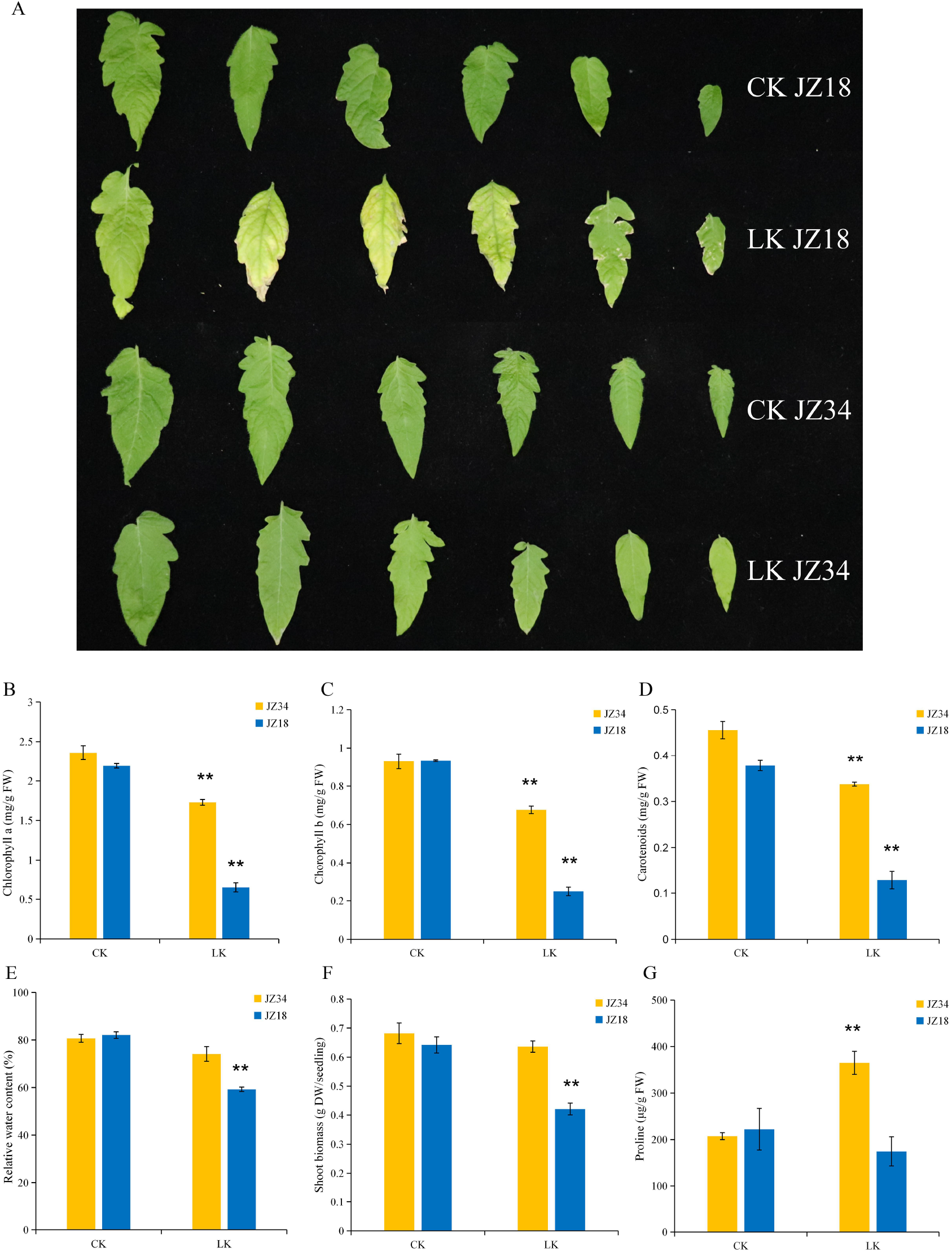
Leaves phenotype of JZ18 and JZ34 plants under control K^+^ (4 mM) and LK (0.5 mM) stress conditions for 21 days. (A) Leaves phenotype, (B) Chlorophyll a content, (C) Chlorophyll b content, (D) Carotenoids content, (E) Relative water content, (F) Shoot biomass and (G) Proline content in JZ18 and JZ34 plants under LK and control conditions for 21 days. The presented data are the means ± SE of three independent experiments (n=12). Asterisks indicate significant difference between JZ18 and JZ34 plants (*P < 0.05, **P < 0.01).

### 3.2 Na^+^, K^+^, Ca^2+^ and Mg^2+^ content in roots and leaves of JZ18 and JZ34 plants

We examined the content of K^+^, Na^+^, Ca^2+^ and Mg^2+^ under LK conditions for 0 days, 1 days, 3 days and 7 days in both roots and shoots of JZ18 and JZ34 plants. In response to LK stress, compared with JZ18, JZ34 exhibited higher K^+^ content in both roots and shoots at various stages after LK treatment (**Fig. 3** A-B).Without LK treatment (0 days), there was no difference between JZ18 and JZ34 plants. The Na^+^ content exhibited a opposite trend to the K^+^ content, which was less in JZ34 than JZ18 in both roots and shoots (**Fig. 3** C-D). So the Na^+^/K^+^ ratio in both roots and shoots of the JZ34 was lower than the JZ18 after LK treatment, and this difference was most significant in the roots after LK treatment for 7 days (**Fig. 3** E-F). Base on the above, Na^+^/K^+^ homeostasis of JZ18 has been severely damaged in the root after LK treatment for 7 days.

**Fig. 3.**
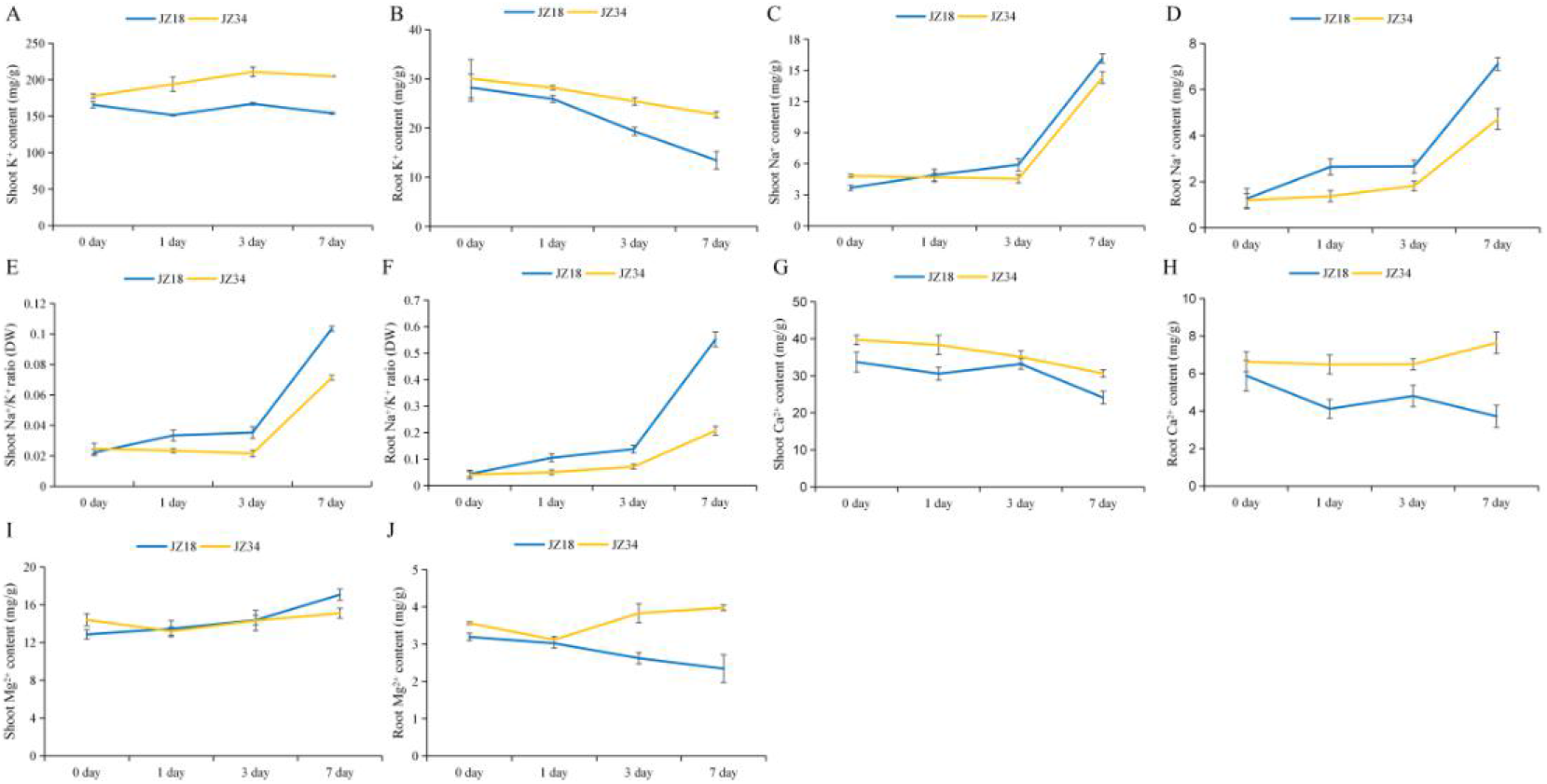
K^+^, Na^+^, Ga^2+^, and Mg^2+^ accumulation in root and shoot tissues of JZ18 and JZ34 plants in response to LK stress. (A) K^+^ content in shoot, (B) K^+^ content in root, (C) Na^+^ content in shoot, (D) Na^+^ content in root, (E) Na^+^/K^+^ ratio in shoot, (F) Na^+^/K^+^ ratio in root, (G) Ca^2+^ content in shoot, (H) Ca^2+^ content in root, (I) Mg^2+^ activity in shoot, and (J) Mg^2+^ activity in root of JZ18 and JZ34 plants under LK stress conditions for 0 days, 1 days, 3 days, and 7 days. The presented data are the means ± SE of three independent experiments (n=12).

The accumulation of Ca^2+^ and Mg^2+^ has been reported to be associated with LK response (Behera *et al.* 2016; Kocourková *et al.* 2020; Dong *et al.* 2021), and we measured the content of Ca^2+^ and Mg^2+^ in both JZ18 and JZ34 plants after LK treatment. The content of Ca^2+^ and Mg^2+^ were higher in the root of JZ34 than JZ18,while there were no significant difference in the shoots (**Fig. 3** G-J). The results implies that JZ34 may relieve the LK stress through the Ca^2+^ and Mg^2+^ signaling pathway (Wang *et al.* 2021).

### 3.3 ROS accumulation and antioxidative competence in JZ18 and JZ34 plants under LK stress

Normally, plants will produce a large amount of reactive oxygen species (ROS) under LK stress (Mittler 2002; R and Schachtman 2004). In both JZ18 and JZ34 plants, the ROS content exhibited an increase trend at the onset of the treatment of LK stress and then declining at late treatment stages, suggesting that the oxidative damage caused by LK stress occurs in the early of LK stress (**Fig. 4** A-D). In general, after treatment with LK, the content of O_2^−^_ and H_2_O_2_ in JZ18 is higher than JZ34, no matter in the root or shoot, which implies that the JZ34 accumulated less ROS in comparison with JZ18. MDA as an index of cellular damage in response to LK stress. After LK stress treatment, the MDA content in JZ34 roots was lower than JZ18 (**Fig. 4** E-F). These results suggest that the lipid peroxidation was lower and the membrane stability was higher in JZ34 roots than JZ18 after LK treatment.

**Fig. 4.**
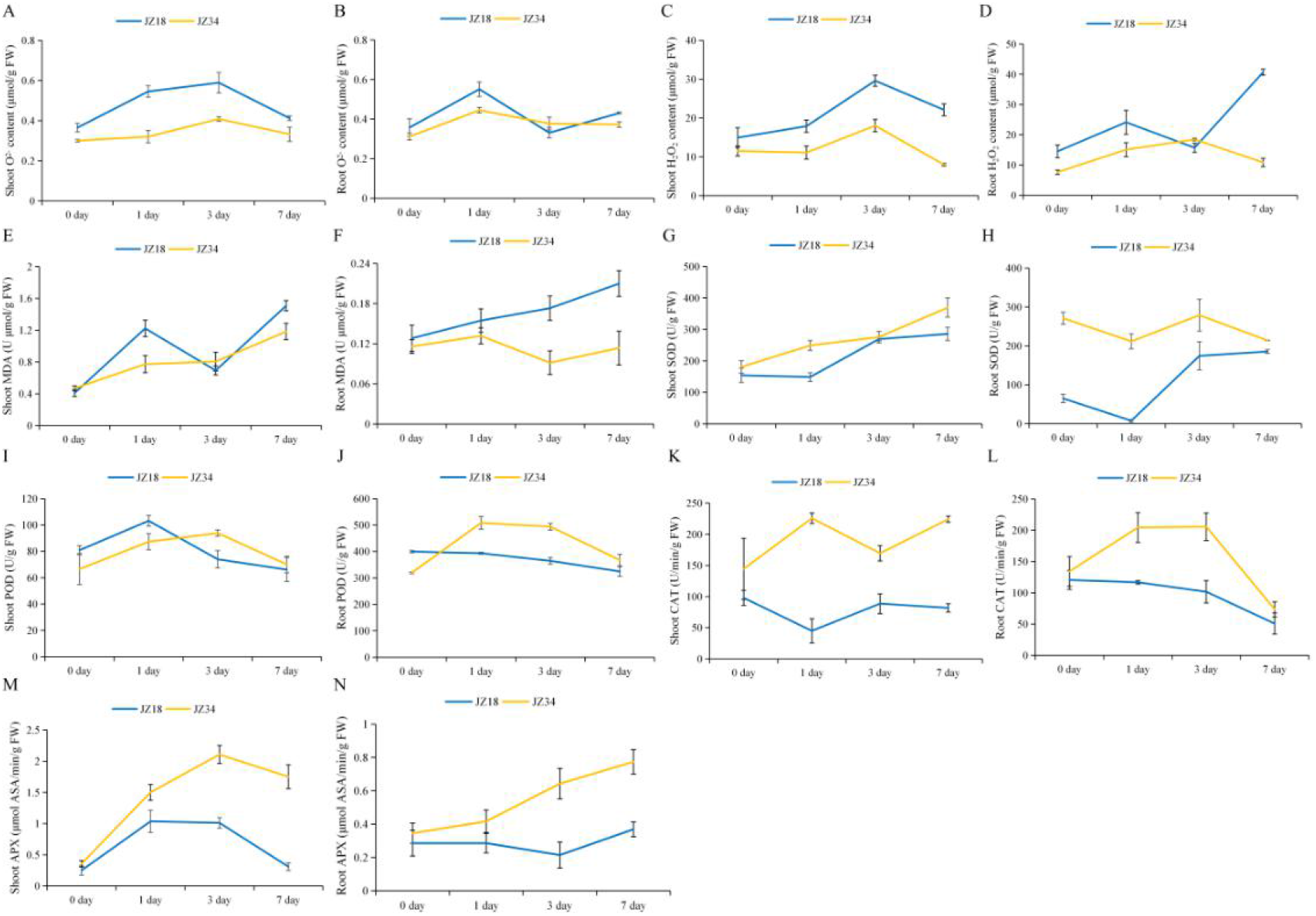
The responses of ROS accumulation and antioxidative competence in JZ18 and JZ34 plants to LK stress. (A) O^2−^ content in shoot (B) O^2−^ content in root (C) H_2_O_2_ content in shoot (D) H_2_O_2_ content in root (E) MDA content in shoot (F) MDA content in root (G) SOD activity in shoot (H) SOD activity in root (I)POD activity in shoot (J) POD activity in root (K) CAT activity in shoot (L) CAT activity in root (M) APX activity in shoot (N) APX activity in root of JZ18 and JZ34 plants under LK conditions for 0 days, 1 days, 3 days and 7 days. The presented data are the means ± SE of three independent experiments (n=12).

The activity of antioxidative enzymes, including superoxide dismutase (SOD), peroxidase (POD), catalase (CAT), and ascorbate peroxidase (APX) in both JZ18 and JZ34 plants was determined at various time. With the LK treatment time, the activity of antioxidative enzyme has changes differently to respond LK stress (**Fig. 4** G-N). The activity of SOD and POD showed no significant differences between JZ18 and JZ34 in the shoots after LK treatment, while the activity of SOD and POD was higher in the roots of JZ34 than JZ18 after LK treatment (**Fig. 4** G-J). CAT and APX showed the same trends, that is, the enzyme activity were significantly higher in both roots and shoots of JZ34 plants compared to JZ18 plants after LK treatment (**Fig. 4** K-N).

All the results demonstrate that in the roots and shoots, JZ34 has lower ROS accumulation and less lipid peroxidation and higher the activity of antioxidative enzymes in comparison with JZ18 in most of the periods under LK stress.

### 3.4 Inheritance analysis of LK tolerance in F_2_ generation

To investigate the genetic basis of LK tolerance, we measured the F_2_ individuals corresponding parameters to confirm the relationship between root length and K^+^ content. The Pearson correlation coefficients (R^2^) between root length and K^+^ content were calculated. There was a significant positive correlation between the two parameters (R^2^ =0.72, P < 0.01) suggesting that root length in tomato is indeed a reliable indicator for K^+^ content (**Fig. 5**).

**Fig. 5.**
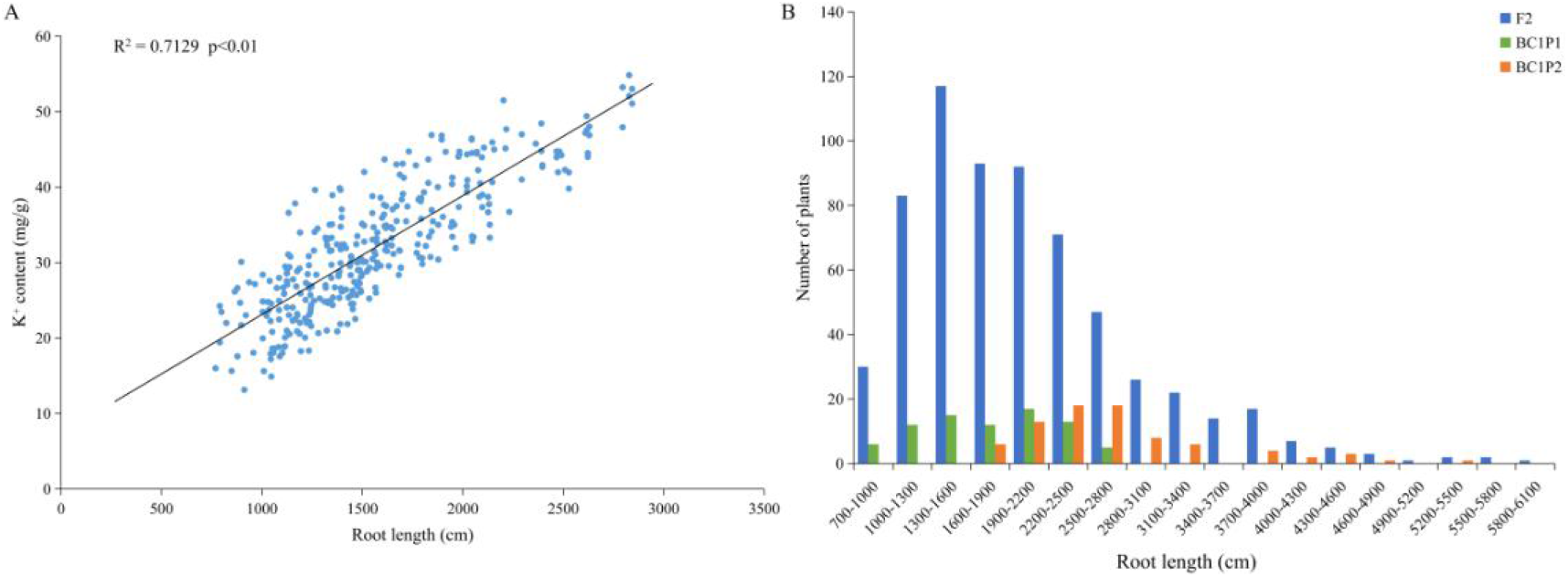
(A) Correlation of the root length and K^+^ content. (B) Frequency distributions of the lateral root length in BC_1_P_1_, BC_1_P_2_ and F2 generations.

Under LK conditions, the root length values of six generations (P_1_, P_2_, F_1_, BC_1_P_1_, BC_1_P_2_ and F_2_) are shown in **Table 1**. Compared with the mean of phenotypic values of two parents, the values of F_1_ population were between the two parents for root length. The coefficient of variation of BC_1_P_1_, BC_1_P_2_ and F_2_ showed a higher level than parents in the lateral root length. The coefficient of variations were higher for segregating populations than parents populations, indicating that the segregating generations showed larger genotypic variation. The frequency distribution of root length in BC_1_P_1_, BC_1_P_2_ and F_2_ populations under LK conditions were observed in **Fig. 5**. The results performed clear skewed distribution in backcross populations and normal distribution in F_2_ generation. Thus, the LK resistance was a quantitative trait, which may be controlled by a mixed major plus polygene model.

**Table 1.** Descriptive statistics of the lateral root length in six generations.

| Trait | Generation | Max | Min | Mean | SD | Variance | CV(%) | Skewness | Kurtosis |
| --- | --- | --- | --- | --- | --- | --- | --- | --- | --- |
| Root length/cm | P1 | 1960.07 | 1450.91 | 1726.69 | 140.00 | 19600.71 | 8.11 | -0.28 | -0.85 |
|  | P2 | 3267.98 | 2206.61 | 2657.63 | 334.57 | 111933.92 | 12.59 | 0.32 | -1.11 |
|  | F1 | 3174.75 | 1226.69 | 2376.12 | 637.59 | 406523.98 | 26.83 | -0.43 | -1.21 |
|  | BC1P1 | 7718.74 | 700.57 | 2162.68 | 995.74 | 991500.36 | 46.04 | 2.01 | 8.25 |
|  | BC2P2 | 6843.39 | 708.60 | 2224.82 | 1042.83 | 1087504.72 | 46.87 | 1.60 | 4.11 |
|  | F2 | 6033.68 | 723.13 | 2045.84 | 863.82 | 746192.94 | 42.22 | 1.31 | 2.23 |

### 3.5 The best-fit genetic model and effect analysis

To evaluate a genetic model for root length under K^+^ deficiency, the AIC values and Log Max likelihood values of 24 genetic models were calculated using joint segregation analysis methods in six generations (**Table 2**). Two genetic models with minimum AIC values would be identified as candidate models of root length. The AIC values of the E-0 and E-1 models were 16025.5334 and 16018.3735, respectively. Next, the goodness-of-fit test was implemented for candidate models to select the optimal model via Uniformity test (U_2_^1^, U_2_^2^ and U_2_^3^), Smirnov test (nW^2^) and Kolmogorov test (D_n_) (**Table 3**). The genetic model with the least number of values achieving statistically significant was identified as the best-fit model. The number of values reaching the significance level for the E-0 and E-1 models were both 1. It’s sensible for models selection to combine the AIC values with the goodness-of-fit test. Therefore, the E-1 (two additive-dominance-epistasis major gene plus additive-dominance polygenes) was the best-fit model for the root length under LK stress conditions. These results suggested that the two major gene plus polygenes regulated the inheritance model of LK resistance.

**Table 2.**
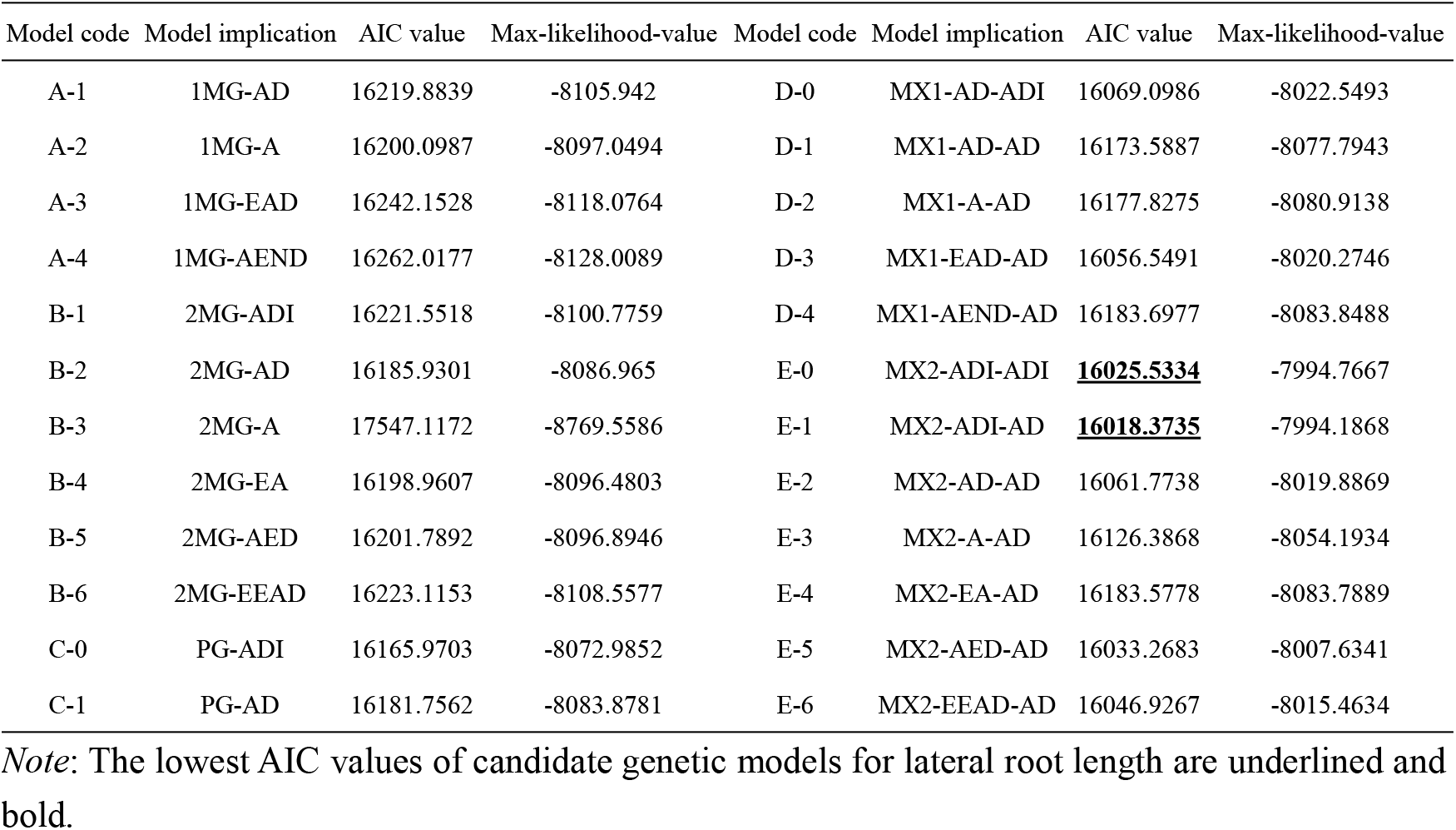
Akaike’s Information Criterion (AIC) value under various genetic models for the lateral root length

| Model code | Model implication | AIC value | Max-likelihood-value | Model code | Model implication | AIC value | Max-likelihood-value |
| --- | --- | --- | --- | --- | --- | --- | --- |
| A-1 | 1MG-AD | 16219.8839 | -8105.942 | D-0 | MX1-AD-ADI | 16069.0986 | -8022.5493 |
| A-2 | 1MG-A | 16200.0987 | -8097.0494 | D-1 | MX1-AD-AD | 16173.5887 | -8077.7943 |
| A-3 | 1MG-EAD | 16242.1528 | -8118.0764 | D-2 | MX1-A-AD | 16177.8275 | -8080.9138 |
| A-4 | 1MG-AEND | 16262.0177 | -8128.0089 | D-3 | MX1-EAD-AD | 16056.5491 | -8020.2746 |
| B-1 | 2MG-ADI | 16221.5518 | -8100.7759 | D-4 | MX1-AEND-AD | 16183.6977 | -8083.8488 |
| B-2 | 2MG-AD | 16185.9301 | -8086.965 | E-0 | MX2-ADI-ADI | **<u>16025.5334</u>** | -7994.7667 |
| B-3 | 2MG-A | 17547.1172 | -8769.5586 | E-1 | MX2-ADI-AD | **<u>16018.3735</u>** | -7994.1868 |
| B-4 | 2MG-EA | 16198.9607 | -8096.4803 | E-2 | MX2-AD-AD | 16061.7738 | -8019.8869 |
| B-5 | 2MG-AED | 16201.7892 | -8096.8946 | E-3 | MX2-A-AD | 16126.3868 | -8054.1934 |
| B-6 | 2MG-EEAD | 16223.1153 | -8108.5577 | E-4 | MX2-EA-AD | 16183.5778 | -8083.7889 |
| C-0 | PG-ADI | 16165.9703 | -8072.9852 | E-5 | MX2-AED-AD | 16033.2683 | -8007.6341 |
| C-1 | PG-AD | 16181.7562 | -8083.8781 | E-6 | MX2-EEAD-AD | 16046.9267 | -8015.4634 |

*Note:* The lowest AIC values of candidate genetic models for lateral root length are underlined and bold.

**Table 3.**
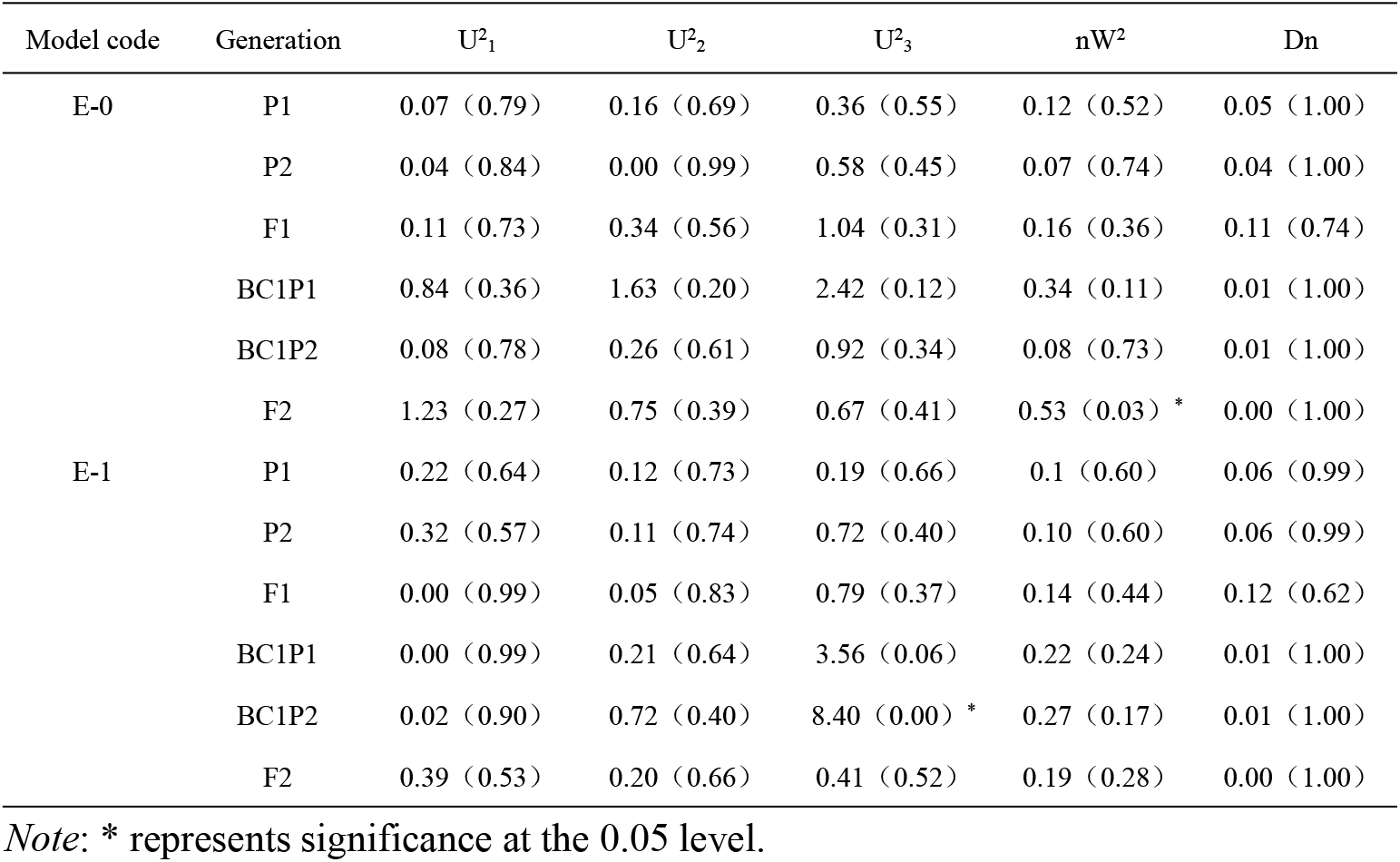
Test for goodness-of-fit of the candidate genetic models

The first-order genetic and second-order genetic parameters of the optimal inheritance models for the root length under LK stress were listed in **Table 4**. Root length exhibited equal additive effect in two major gene due to d_a_=d_b_. The dominance effect of the first major gene were greater than those of the second major gene for root length. The potential ratio |h_a_/d_a_| and |h_b_/d_b_| of the major gene were less than 1, suggesting that the dominance effect was smaller than the additive effect of two major genes. The additive plus dominance interaction effect (j_ab_) and dominance plus additive interaction effect (j_ba_) were positive values, indicating that these interactions between two major genes improved root length to enhance LK tolerance. Thus, additive, dominant and epistatic effects are important for the inheritance of tomato LK resistance.

**Table 4.** The estimate of genetic parameters of the best-fit model for the six traits

| 1st order parameter | Estimate | 2nd order parameter | Estimate |  |  |
| --- | --- | --- | --- | --- | --- |
| | E-1 | | $BC_1P_1$ | $BC_1P_2$ | $F_2$ |
| $d_a$ | -692.85 | $\sigma_p^2$ | 991500.36 | 1087504.72 | 746192.94 |
| $d_b$ | -692.85 | $\sigma_{mg}^2$ | 8821.42 | 552148.08 | 519050.12 |
| $h_a$ | -208.82 | $\sigma_{pg}^2$ | 823470.43 | 374195.43 | 59394.40 |
| $h_b$ | 76.37 | $h_{mg}^2$ | 0.88 | 50.41 | 69.45 |
| $i$ | 612.15 | $h_{pg}^2$ | 82.25 | 34.16 | 7.95 |
| $l$ | -375.86 | | | | |
| $j_{ab}$ | 144.72 | | | | |
| $j_{ba}$ | 429.90 | | | | |
| $h_a/d_a$ | 0.30 | | | | |
| $h_b/d_b$ | -0.11 | | | | |

*Note:* $d_a$ : additive effect of the first pair major gene; $d_b$ : additive effect of the second pair major gene; $h_a$ : dominant effect of the first pair major gene; $h_b$ : dominant effect of the second pair major gene; $i$ : additive effect plus additive effect of the two major genes; $l$ : dominant effect plus dominant effect of the two major genes; $j_{ab}$ : additive effect plus dominant effect of the two major genes; $j_{ba}$ : dominant effect plus additive effect of the two major genes; $h_a/d_a$ : dominance degree of the first major gene; $h_b/d_b$ : dominance degree of the second major gene; $\sigma_p^2$ : phenotypic variance; $\sigma_{mg}^2$ : major gene variance; $\sigma_{pg}^2$ : polygene variance; $h_{mg}^2(\%)$ : major gene heritability; $h_{pg}^2(\%)$ : polygene heritability.

The heritability of major gene from the BC_1_P_1_, BC_1_P_2_ and F_2_ populations were 0.88%, 50.4% and 69.45%, respectively, indicating the diversity of genetic inheritance. The heritability of major gene were greater than the polygene heritability for F_2_ generation, suggesting that LK stress was primarily regulated bygenetic factors. The root length were slightly affected by environmental factors due to the high heritabilit, and indicating that selection for root length in early generations is most efficient.

### 3.6 QTL-seq analysis combining SNP-index and InDel-index

To identify the QTL for LK tolerance, we compared two extreme pools from the F_2_ population, a LK resistance pool (R-pool) and LK susceptibility (S-pool) using BSA-seq. A total of 118.13 Gb valid data were obtained by Illumina sequencing, including 32.63 Gb from the R-pool and 39.79 Gb from the S-pool, all of high quality (91.38% > Q30 > 93.82%) and with a stable GC content (41.00% > GC > 46.83%) (**Table 5**). We used Venn diagrams to demonstrate the relationships between SNPs and InDels among the parents and the mixed pools (**Fig. 6** A-B). A total of identified 990,251 SNPs and 217,061 InDels in the four pools in comparison with the reference genome respectively. These high-quality data lay a solid foundation for subsequent analysis.

**Table 5.** Sequencing data quality statistics

| Sample | Raw_Reads | Raw_Bases | Valid_Reads | Valid_Bases | Valid% | Q20% | Q30% | GC% |
| --- | --- | --- | --- | --- | --- | --- | --- | --- |
| JZ18 | 164366004 | 24.65G | 155226540 | 23.28G | 94.44 | 96.74 | 91.63 | 43.95 |
| JZ34 | 159970176 | 24.00G | 149552510 | 22.43G | 93.49 | 96.59 | 91.38 | 41.00 |
| S_F2 | 273786410 | 41.07G | 265248210 | 39.79G | 96.88 | 97.86 | 93.82 | 44.17 |
| R_F2 | 242359902 | 36.35G | 217517696 | 32.63G | 89.75 | 96.77 | 92.14 | 46.83 |

Note: Q20 is the proportion of bases with quality value $\geq 20$ (base recognition accuracy rate $> 99\%$ ); Q30 is the proportion of bases with quality value $\geq 30$ (base recognition accuracy rate $> 99.9\%$ ); GC(%) is the content of bases G and C and the proportion of total bases.

**Fig. 6.**
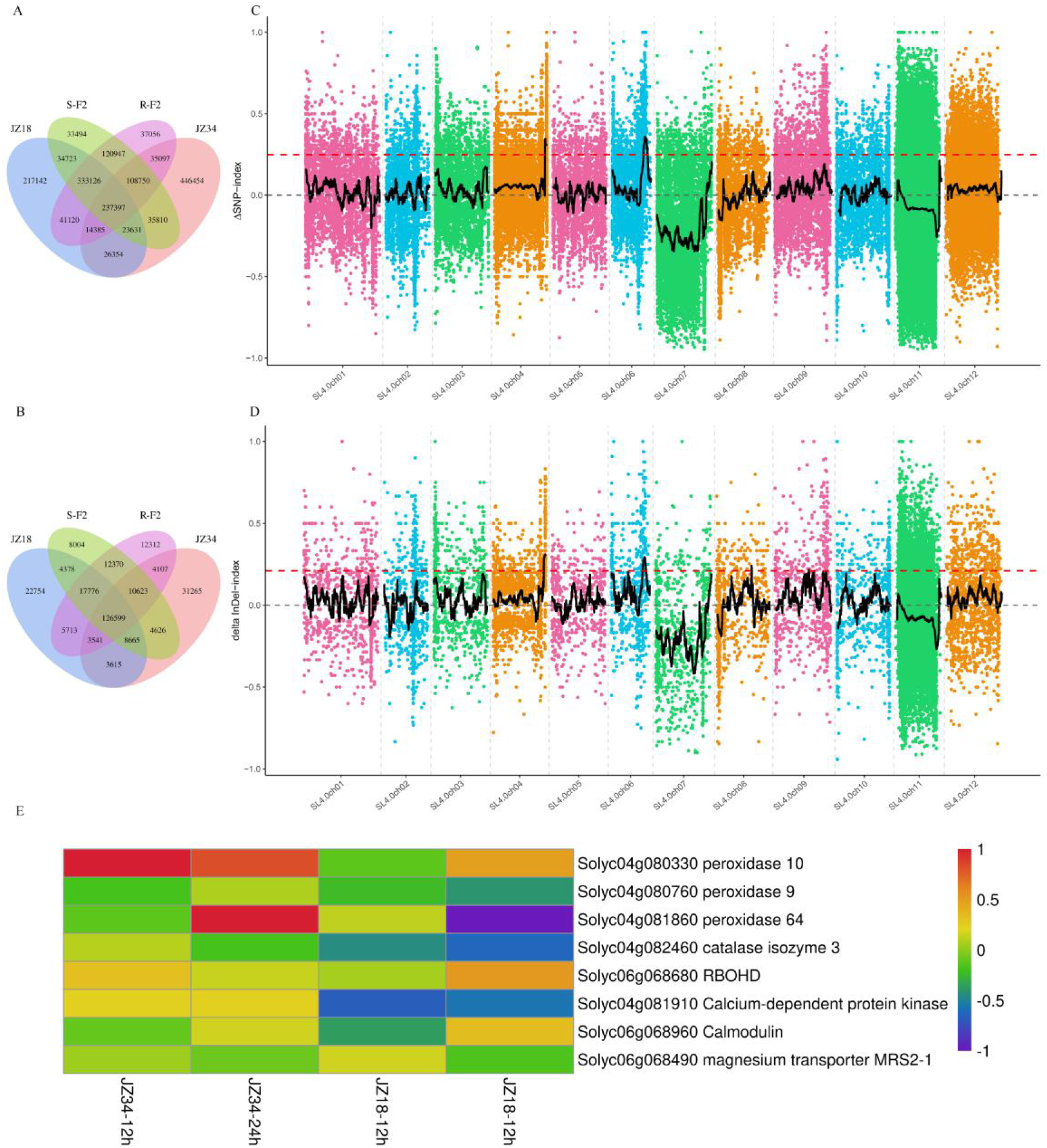
SNP statistics and BSA analysis. (A) Venn diagram of SNP in the four pools. (B) Venn diagram of InDel in the four pools. (C) SNP-index algorithm to map root length based LK stress gene. (D) InDel-index algorithm to map root length based LK stress gene. (E) Differentially expressed genes involved ROS and ion signaling pathways from our previous report (Zhao *et al.* 2018).

To detect the major QTLs responsible for LK tolerance, we used SNP-index and InDel-index association algorithms (**Fig. 6** C-D). As shown in Fig, 61.87-64.45 Mb (2.58 Mb) region on chromosome A04 and 39.27-40.65 (1.38Mb) region on chromosome A06 exhibiting significant linkage were identified as the candidate region, and both the two different methods mapped these QTL at a 95% significance level. The result is consistent with the analysis result of the previous genetic model that LK resistance was controlled by two pairs of major genes. Therefore, the two candidate region were selected as the major QTL for LK resistance. On chromosome A04 candidate region annotated a total of 369 genes, including 18 non-synonymous genes and 5 frameshifted genes. On chromosome A06 candidate region annotated a total of 198 genes, including 9 non-synonymous genes and 3 frameshifted genes.

The Physiological results were used to identify the candidate genes within the 2.58 Mb and 1.38 Mb intervals. A total of 4 genes were linked to antioxidant, inclding *Solyc04g080330* (peroxidase 10), *Solyc04g080760* (peroxidase 9), *Solyc04g081860* (peroxidase 64) and *Solyc04g082460* (catalase isozyme 3). *Solyc06g068680* (RBOHD) was responsible for the generation of LK-induced ROS signals. In addition, *Solyc04g081910* (Calcium-dependent protein kinase) and *Solyc06g068960* (Calmodulin) transferred Ca^2+^ signal, and *Solyc06g068490* (magnesium transporter MRS2-1) involved in Mg^2+^ transporter. The transcript levels of these genes were significantly changed under LK conditions in JZ18 and JZ34 (**Fig. 6** E). In JZ18 plant, the expression levels of most genes were down-regulated with LK treatment. However, most genes in JZ34 plant were up-regulated, especially *Solyc04g080330* and *Solyc04g081860*.

## 4. Discussion

### 4.1 Root length is an important morphologically adaptive traits for LK

In our study, long-term LK treament, the JZ34 plants maintained normal root growth and kept leaves green. However, in JZ18 plants, short-term LK treatment caused damage to normal root growth, and after further increasing the time of LK treatment, the leaves of JZ18 plants gradually showed turned yellow-green color. The K^+^ uptake by plant root cells as well as K^+^ transport inside plants are conducted by a large number of K^+^ channels and transporters (Wang and Wu 2013; Very *et al.* 2014). K^+^ deficiency enhances the elongation of root hair and inhibites primary root growth in *Arabidopsis thaliana* (Jung *et al.* 2009; Qin *et al.* 2019).Thus, root characterization of JZ34 plant under LK stress is a direct evidence to prove its LK resistance. Moreover, we measured some physiological index to support the results of phenotypic observation. The JZ34 had higher root biomass, root length, root area and root fork, which exhibited the strong root growth ability of JZ34 plant under LK conditions. Under LK stress, the biomass, chlorophyll content, RWC and proline content in the leaves were higher in JZ34 plants than the JZ18 plants, which may be owing to the JZ34 plants adapted to LK stress conditions through maintaining normal root growth. These findings showed that JZ34 is a LK resistance tomato variety through normal growth of root to absorb more K^+^.

The elongation of root length is an essential adaptive trait for LK tolerance. The change of K^+^ concentration influences root developmental processes, including primary root growth, lateral root formation and root-hair formation (López-Bucio *et al.* 2003). The early investigation to screen K^+^ high efficient tomato varieties under K^+^ deficiency, the root activity was used as K^+^ efficiency genotype screening optimal index (Yang *et al.* 2015). In addition, several LK resistance genes have been mapped through the observation of the growth of root length under K^+^ deficiency in *Arabidopsis*, including *CIPK23*, *AtKC1*, *MYB59* and *NPF7.3/NRT1.5* (Xu *et al.* 2006; Wang *et al.* 2010; Li *et al.* 2017; Du *et al.* 2019), indicating the great contribution of the root length for the genetic investigation of LK stress. There was a significant difference for root length between the LK-tolerance line JZ34 and LK-sensitive line JZ18. The highly significant positive correlation between root length and K^+^ content showed the root length is well indicator of LK resistance. Due to counting root length is more simple and more direct than measuring of K^+^ content, root length offers an easier choice for assessing LK tolerance.

### 4.2 LK resistance is a quantitative trait controlled by multiple genes in tomato

Improving of LK tolerance is the one of important targets in tomato breeding. The genetic analysis of tolerance to LK in maize have been also estimated using joint segregation analysis, showing that tolerance to LK stress in maize was dominated by one major gene plus polygene (Li *et al.* 2011). However, the studies of genetic inheritance to LK stress in tomato were rarely applied breeding. Here, we attempted to further clarify the inheritance mechanism of LK resistance in tomato, six generations (P_1_, P_2_, F_1_, B_1_, B_2_ and F_2_) were used to analyze the major gene plus polygene inheritance model based root length under LK stress. Root length under LK conditions in tomato was a quantitative trait, which were controlled by a group of genes with different effects. Under LK condition, the inheritance model and inheritance effect of root length was assessed by joint segregation analysis based on the AIC values, max-likelihood value, and goodness-of-fit test. The results suggested that the best-fit model was regulated by two additive-dominance-epistasis major genes plus additive-dominance polygene inheritance model (E-1). In early study, the inheritance methods of K^+^ utilization efficiency have been similarly considered as additive-dominance-epistasis polygene genetic model (Gabelman and Loughman 1987). In our study, the major gene heritability of BC_1_P_1_, BC_1_P_2_ and F_2_ populations were 0.88%, 50.4% and 69.45%, respectively, indicating that the inheritance method of each populations was diverse due to the genetic background of the parents’ traits (Ye *et al.* 2017). The phenotype characters were influenced by close integration between genetic effect and environmental influence. Our genetic analysis results showed that the root length was affected by smaller environmental influence, and the trait choice should occur in early populations. Further SNP-index and InDel-index linkage analysis with two extreme mixed pool of root length contains approximately 30 individuals generally maps the two target regions. The major QTL interval were 2.38Mb at the end of chromosome 4 and 1.38 Mb at the chromosome 6, which are consistent with previous analysis of the genetic model controlled by two major gene.

### 4.3 Ion and antioxidant signal may be involved in the mechanism of LK stress

The ability to maintain ion homeostasis play an essential role for LK resistance and involves a network of transport processes that regulates uptake, extrusion through the plasma membrane in plants (Apse and Blumwald 2007). In our study, the K^+^ content was higher in JZ34 compared with JZ18 plants after LK treatment, suggesting that the activity of K^+^ transporter or K^+^ channel might be improved in JZ34 plants to respond LK stress. By contrast, in root and shoot tissues, the JZ34 showed lower Na^+^ content than the JZ18 plants, which caused a higher Na^+^/K^+^ ratio in JZ18 plants after LK stress. Especially, after LK treatment for 7 days in the roots, these Na^+^/K^+^ ratio existed a significant difference between JZ34 and JZ18 plants, showing that the damage of ion homeostasis becomes more serious as the treatment time of LK stress increases in the roots of JZ18 plants.

Recent research found that high concentrations of Mg^2+^ disrupt K^+^ homeostasis, and that transcription of K^+^ homeostasis-related genes CIPK9 and HAK5 is changed to limit the elongation of root length.(Kocourková *et al.* 2020). Ca^2+^ signal also can be triggered rapid K^+^ deprivation in the root, in which Ca^2+^ induces CIF peptides to activate SGN3-LKS4/SGN1 receptor complexes, and then convey HAK5 K^+^ transporter induction (Wang *et al.* 2021). Our characterization of the two candidate regions at the chromosome 4 and 6, existed 3 genes, *Solyc04g081910* (Calcium-dependent protein kinase), *Solyc06g068960* (Calmodulin) transferred Ca^2+^ signal, and *Solyc06g068490* (magnesium transporter MRS2-1) involved in Mg^2+^ transporter. Especially, *Solyc04g081910* (Calcium-dependent protein kinase) gene expression was down-regulated in JZ18 plants. Thus, the results obtained in root and shoot tissues, the lower accumulation of Ca^2+^ and Mg^2+^ in JZ18 plants than JZ34 plants suggested JZ34 may respond to LK stress through maintain normal Ca^2+^ and Mg^2+^ signal.

K^+^ deficiency induces the accumulation of ROS and generates ROS associated injury (Mittler 2002; R and Schachtman 2004). LK stress inducing the production of ROS for JZ18 and JZ34 plants was found in previous study, but the reason have remained unknown (Zhao *et al.* 2018). In the present study, in roots and shoots, LK stress led to ROS accumulation and increased MDA content in JZ18, which further resulted in membrane lipid peroxidation and cell membrane damage. The JZ34 plants exhibited slight ROS accumulation and cell membrane damage. Moreover, JZ34 treated by LK stress had a higher proline levels than normal K^+^ treatment, which protect cells against increased ROS levels. To neutralize the injury of oxidative stress, plants use precisely controlled ROS scavenging strategies, such as enzymatic systems (Mittler 2002; Golldack *et al.* 2014). Interestingly, 4 genes related to antioxidant enzymes were selected in candidate regions, including *Solyc04g080330* (peroxidase10), *Solyc04g080760* (peroxidase9), *Solyc04g081860* (peroxidase64), and *Solyc04g082460* (catalase isozyme3). The expression of these gene were up-regulated under LK conditions in JZ34 plants, while in JZ18 plants were down-regulated. After LK treatment, in roots and shoots, JZ34 had higher activities of antioxidant enzymes for SOD, APX, and CAT than JZ18 plants at all periods, implying that the JZ34 plants might be involved to the improved activities of antioxidant enzymes to reduce the injury of oxidative stress, and then enhanced LK resistance. In the recent study showed that plants sense K^+^ deficiency and trigger rapid K^+^ and Ca^2+^ signals, and then phosphorylates and activates RBOHC/D/F for ROS signal formation to convey HAK5 K^+^ transporter induction (Wang *et al.* 2021). In the candidate regions, we just found *Solyc06g068680* (RBOHD) can induced the accumulation of ROS, and its expression was up-regulated in JZ18 and JZ34 plants under LK conditions. This also indicated that, LK stress led to the decrease of K^+^, Ca^2+^, and Mg^2+^ content, which may translate signals to enhance ROS accumulation, and further regulated the expression of genes related to LK in response to LK stress in tomato.

In conclusion, JZ34 improved Na^+^/K^+^ homeostasis, and repressed ROS accumulation under LK stress, suggesting that JZ34 can enhance the LK stress tolerance compared with JZ18 plants. The method of major gene plus polygene model with the application of the joint segregation analysis, we showed that the root length trait under LK stress might regulated by two additive-dominance-epistasis major genes plus additive-dominance polygene inheritance model (E-1). Through BSA-seq, two major-effect QTLs that were responsible for the phenotypic variation of root length in tomato under LK stress condition were identified. Combine with physiological and mapping results of LK stress responses in JZ18 and JZ34 plants enabled us select several interesting candidate genes controlling the LK tolerance. These results will provide some instructions for fine mapping and breeding of LK resistance in the future, and laid the theoretical basis for the mining and screening of LK resistance tomato resources.

## Acknowledgements

This research was supported by the National Key Research and Development Program of China (2019YFD100030) and the National Natural Science Founding of China (No.31372054).

## Data availability

The data underlying this article are available in the article and in its online supplementary material.

## References

Apse M. P., and E. Blumwald, 2007 Na^+^ transport in plants. Febs Lett. 581: 2247–2254. https://doi.org/10.1016/j.febslet.2007.04.014

Asins M. J., I. Villalta, M. M. Aly, R. Olías, P. Álvarez De Morales, et al., 2013 Two closely linked tomato HKT coding genes are positional candidates for the major tomato QTL involved in Na^+^/K^+^ homeostasis. Plant, Cell Environ. 36: 1171–1191. https://doi.org/10.1111/pce.12051

Behera S., L. Yu, I. SchmitzTHom, X. Wang, and W. Yi, 2016 Two spatially and temporally distinct Ca^2+^ signals convey Arabidopsis thaliana responses to K^+^ deficiency. New Phytol. 213: 739. https://doi.org/10.1111/nph.14145

Cao Y., A. D. M. Glass, and N. M. Crawford, 1993 Ammonium inhibition of Arabidopsis root growth can be reversed by potassium and by auxin resistance mutations *aux1*, *axr1*, and *axr2*. Plant Physiol. 102: 983–989.

Ding Y., W. Luo, and G. Xu, 2010 Characterisation of magnesium nutrition and interaction of magnesium and potassium in rice. Ann. Appl. Biol. 149: 111–123. https://doi.org/10.1111/j.1744-7348.2006.00080.x

Dong Q., B. Bai, B. O. Almutairi, and J. Kudla, 2021 Emerging roles of the CBL-CIPK calcium signaling network as key regulatory hub in plant nutrition. J. Plant Physiol. 257: 153335. https://doi.org/10.1016/j.jplph.2020.153335

Du X. Q., F. L. Wang, H. Li, S. Jing, M. Yu, et al., 2019 The transcription factor MYB59 regulates K^+^/NO_3^−^_ translocation in the arabidopsis response to low K^+^ stress. Plant Cell 31: 699–714. https://doi.org/10.1105/tpc.18.00674

Fageria V. D., 2001 Nutrient interactions in crop plants. J. Plant Nutr. 24: 1269–1290. https://doi.org/10.1081/PLN-100106981

Fang Y., W. Wu, X. Zhang, H. Jiang, W. Lu, et al., 2015 Identification of quantitative trait loci associated with tolerance to low potassium and related ions concentrations at seedling stage in rice (Oryza sativa L.). Plant Growth Regul. 77: 157–166. https://doi.org/10.1007/s10725-015-0047-9

Gabelman W. H., and B. C. Loughman, 1987 Genetic aspects of plant mineral nutrition. J. Appl. Ecol. 28: 745. https://doi.org/10.1007/978-94-009-2053-8

Gai J. Y., and J. K. Wang, 1998 Identification and estimation of a QTL model and its effects. Theor. Appl. Genet. 97: 1162–1168. https://doi.org/10.1007/s001220051005

Golldack D., C. Li, H. Mohan, and N. Probst, 2014 Tolerance to drought and salt stress in plants: Unraveling the signaling networks. Front Plant 5: 151. https://doi.org/10.3389/fpls.2014.00151

Jung J. Y., R. Shin, and D. P. Schachtman, 2009 Ethylene mediates response and tolerance to potassium deprivation in Arabidopsis. Plant Cell 21: 607–621. https://doi.org/10.1105/tpc.108.063099

Kocourková D., Z. Krčková, P. Pejchar, K. Kroumanová, T. Podmanická, et al., 2020 Phospholipase Dα1 mediates the high-Mg^2+^ stress response partially through regulation of K^+^ homeostasis. Plant Cell Environ. 43: 2460–2475. https://doi.org/10.1111/pce.13831

Kong F. M., Y. Guo, X. Liang, C. H. Wu, Y. Y. Wang, et al., 2013 Potassium (K) effects and QTL mapping for K efficiency traits at seedling and adult stages in wheat. Plant Soil 373: 877–892. https://doi.org/10.1007/s11104-013-1844-4

Koyama M. L., A. Levesley, R. M. Koebner, T. J. Flowers, and A. R. Yeo, 2001 Quantitative trait loci for component physiological traits determining salt tolerance in rice. Plant Physiol. 125: 406–422. https://doi.org/doi:10.1104/pp.125.1.406

Li X. T., M. J. Cao, H. Q. Yu, and X. G. Wang, 2011 Genetic analysis of tolerance to low-potassium stress in maize using mixed model of major gene plus polygene. J. Maize Sci. 54: 235–241 in Chinese with English abstract. https://doi.org/10.13597/j.cnki.maize.science.2011.04.033

Li H., M. Yu, X. Q. Du, Z. F. Wang, W. H. Wu, et al., 2017 NRT1.5/NPF7.3 functions as a proton-coupled H^+^/K^+^ antiporter for K^+^ loading into the xylem in arabidopsis. Plant Cell 29: 2016–2026. https://doi.org/10.1105/tpc.16.00972

Lin H., M. Zhu, M. Yano, J. Gao, Z. Liang, et al., 2004 QTLs for Na^+^ and K^+^ uptake of the shoots and roots controlling rice salt tolerance. Theor. Appl. Genet. 108: 253–260. https://doi.org/10.1007/s00122-003-1421-y

López-Bucio J., A. Cruz-Ram\irez, and L. Herrera-Estrella, 2003 The role of nutrient availability in regulating root architecture. Curr. Opin. Plant Biol. 6: 280–287. https://doi.org/10.1016/S1369-5266(03)00035-9

Mittler R., 2002 Oxidative stress, antioxidants and stress tolerance. Trends Plant Sci. 7: 405–410. https://doi.org/10.1016/S1360-1385(02)02312-9

Mogami J., Y. Fujita, T. Yoshida, Y. Tsukiori, H. Nakagami, et al., 2015 Two distinct families of protein kinases are required for plant growth under high external Mg^2+^ concentrations in Arabidopsis. Plant Physiol. 167: 1039–1057. https://doi.org/10.1104/pp.114.249870

Pandit A., V. Rai, S. Bal, S. Sinha, V. Kumar, et al., 2010 Combining QTL mapping and transcriptome profiling of bulked RILs for identification of functional polymorphism for salt tolerance genes in rice (OryzasativaL.). Mol. Genet. Genomics 284: 121–136. https://doi.org/10.1007/s00438-010-0551-6

Pardo J. M., and F. Rubio, 2011 Na^+^ and K^+^ Transporters in Plant Signaling. Springer Berlin Heidelb. https://doi.org/10.1007/978-3-642-14369-4_3

Qin Y. J., W. H. Wu, and Y. Wang, 2019 ZmHAK5 and ZmHAK1 function in K^+^ uptake and distribution in maize under low K^+^ conditions. J. Integr. Plant Biol. 61: 691–705. https://doi.org/10.1111/jipb.12756

R S., and D. P. Schachtman, 2004 Hydrogen peroxide mediates plant root cell response to nutrient deprivation. Proc. Natl. Acad. Sci. 101: 8827–8832. https://doi.org/0027-8424(2004)101:23<8827:HPMPRC>2.0.TX;2-9

Rodríguez-Navarro A., 2000 Potassium transport in fungi and plants. Biochim. Biophys. Acta 1469: 1–30. https://doi.org/10.1016/S0304-4157(99)00013-1

Senbayram M., R. Gransee, V. Wahle, and H. Thiel, 2015 Role of magnesium fertilisers in agriculture: plant-soil continuum. Crop Pasture Sci. https://doi.org/10.1071/CP15104

Shabala S., and Y. Hariadi, 2005 Effects of magnesium availability on the activity of plasma membrane ion transporters and light-induced responses from broad bean leaf mesophyll. Planta 221: 56–65. https://doi.org/http://ecite.utas.edu.au/26477

Song W., R. Xue, Y. Song, Y. Bi, Z. Liang, et al., 2018 Differential response of first-order lateral root elongation to low potassium involves nitric oxide in two tobacco cultivars. J. Plant Growth Regul. 37: 114–127. https://doi.org/10.1007/s00344-017-9711-9

Tsay Y. F., C. H. Ho, H. Y. Chen, and S. H. Lin, 2011 Integration of nitrogen and potassium signaling. Annu. Rev. Plant Biol. 62: 207. https://doi.org/10.1146/annurev-arplant-042110-103837

Very A. A., M. Nieves-Cordones, M. Daly, I. Khan, C. Fizames, et al., 2014 Molecular biology of K^+^ transport across the plant cell membrane: what do we learn from comparison between plant species? J. Plant Physiol. 171: 748–769. https://doi.org/10.1016/j.jplph.2014.01.011

Wang Y., L. He, H. D. Li, J. Xu, and W. H. Wu, 2010 Potassium channel α-subunit AtKC1 negatively regulates AKT1-mediated K^+^ uptake in Arabidopsis roots under low-K^+^ stress. Cell Res. 20: 826–837. https://doi.org/10.1038/cr.2010.74

Wang Y., and W. H. Wu, 2013 Potassium Transport and signaling in higher plants. Annu. Rev. Plant Biol. 64: 451–476. https://doi.org/10.1146/annurev-arplant-050312-120153

Wang Y., and W. H. Wu, 2017 Regulation of potassium transport and signaling in plants. Curr. Opin. Plant Biol. 39: 123–128. https://doi.org/10.1016/j.pbi.2017.06.006

Wang F. L., Y. L. Tan, L. Wallrad, X. Q. Du, A. Eickelkamp, et al., 2021 A potassium-sensing niche in Arabidopsis roots orchestrates signaling and adaptation responses to maintain nutrient homeostasis. Dev. Cell 56: 781–794.e6. https://doi.org/10.1016/j.devcel.2021.02.027

Xu J., H. D. Li, L. Q. Chen, Y. Wang, L. L. Liu, et al., 2006 A protein kinase, interacting with two calcineurin b-like proteins, regulates K^+^ transporter AKT1 in Arabidopsis. Cell 125: 1347–1360. https://doi.org/10.1016/j.cell.2006.06.011

Yang J. L., X. Y. Xu, and J. F. Li, 2015 Research on screening methods of potassium high efficientcy genotypes in tomato seedling stage. North. Hortic. 12: 40–42 in Chinese with English abstract. https://doi.org/10.11937/bfyy.201512012

Ye Y., J. Wu, L. Feng, Y. Ju, M. Cai, et al., 2017 Heritability and gene effects for plant architecture traits of crape myrtle using major gene plus polygene inheritance analysis. Sci. Hortic. (Amsterdam). 225: 335–342. https://doi.org/10.1016/j.scienta.2017.06.065

Zhao Z., G. Zhang, S. Zhou, Y. Ren, and W. Wang, 2017 The improvement of salt tolerance in transgenic tobacco by overexpression of wheat F-box gene TaFBA1. Plant Sci. 259: 71–85. https://doi.org/10.1016/j.plantsci.2017.03.010

Zhao X., Y. Liu, X. Liu, and J. Jiang, 2018 Comparative transcriptome profiling of two tomato genotypes in response to potassium-deficiency stress. Int. J. Mol. Sci. 19. https://doi.org/10.3390/ijms19082402

